# An SR protein is essential for the recovery of malaria parasites from DNA damage and exposure to artemisinin

**DOI:** 10.1101/2021.01.11.426314

**Authors:** Brajesh Kumar Singh, Manish Goyal, Karina Simantov, Yotam Kaufman, Shiri Eshar, Dzikowski Ron

## Abstract

*Plasmodium falciparum*, the parasite responsible for the deadliest form of human malaria, maintains a complex life cycle with a relatively small number of genes. *Pf*SR1 is an alternative splicing factor that regulates expansion of the *P. falciparum* protein repertoire. To further investigate *Pf*SR1 functions, we set to unveil its interactome. We found that *Pf*SR1 interacts with proteins, which are linked to various processes of RNA metabolism in a stage-dependent manner. These include: chromatin re-modeling, transcription, splicing and translation. Intriguingly, some of the *Pf*SR1 interacting proteins are orthologues of proteins implicated in the DNA damage response. We demonstrate that *Pf*SR1 expression is important for preventing the accumulation of DNA damage in proliferating parasites. In addition, following parasites’ exposure to a source of DNA damage, *Pf*SR1 is recruited to damaged foci where it interacts with the phosphorylated core histone *Pf*H2A, which marks damaged chromatin. Furthermore, *Pf*SR1 expression was found to be essential for the ability of the parasite to activate the DNA repair machinery and recover from DNA damage caused by either irradiation or exposure to artemisinin, the first line anti-malarial drug. These findings unveil a novel role of *Pf*SR1 in protecting *P. falciparum* from DNA damage and artemisinin exposure.

## Introduction

Malaria remains one of the deadliest infectious diseases, killing approximately half a million people each year, primarily young children and pregnant women in sub-Saharan Africa (1). *Plasmodium falciparum* is the protozoan parasite responsible for over 90% of malaria deaths and thus is considered the deadliest form of human malaria. This parasite has a complex life cycle alternating between different biological niches in the human host and the Anopheline vector. Surprisingly, *Plasmodium* parasites have evolved this complex biology with approximately 5700 protein coding genes in their genome, which is even lower than the gene number of the budding yeast, *Saccharomyces cerevisiae* that has a much simpler life cycle (2). One way by which eukaryotes can expand their protein repertoire out of a limited gene number is through alternative splicing (AS) of their pre-mRNA transcripts. Alternative splicing is thought to be regulated by splicing factors that recognize specific splicing signals such as enhancers and silencers (ESE and ESS respectively) on pre-mRNA molecules. Serine/arginine-rich (SR) proteins are known splicing factors that bind splicing enhancers in a sequence specific manner. These proteins bind RNA through domains known as recognition motifs (RRM) and contain a distinctive domain enriched in serine and arginine (RS) that is involved in protein-protein interactions and cellular localization (3). Several putative SR proteins were annotated in the *P. falciparum* genome but only *Pf*SR1 (PF3D7_0517300) was shown to function as an alternative splicing factor *in vivo*(4). *Pf*SR1 has two RRM domains and one RS domain which is essential for its nuclear translocation (4). In addition to alternative splicing, *Pf*SR1 regulates mRNA levels by binding to specific RNA motifs (5). In recent years it became clear that RNA processing is linked to the DNA damage response (DDR), and several splicing factors were implicated as gatekeepers of genome stability (6,7). In any living cell, DNA breaks are continuously generated due to replication errors, and the blood stage of malaria parasites that replicate by consecutive mitoses during schizogony are particularly prone to such errors. In addition, their exposure to immune attack by substances that cause oxidative stress could lead to DNA damage (8). Indeed, it was recently suggested that artemisinin, which is used as the first line antimalarial chemotherapeutic, also functions as a DNA damaging agent (9). Interestingly, *Plasmodium* parasites utilize both homologous recombination (HR) and alternative non homologous end joining (A-NHEJ) to repair double strand breaks (DSB) (10,11). Additionally, recent *in silico* analysis identified components of the DNA mismatch repair pathway in the *P. falciparum* genome (12), which have been previously proposed to play a role in generating diversity and drug resistance in malaria parasites (13–15). Here we show that *Pf*SR1 interacts with proteins that play a role in different processes of RNA metabolism as well as with proteins involved in DNA damage response. We demonstrate that *Pf*SR1 is recruited to the site of DNA damage where it interacts with the phosphorylated core histone PfH2A (γ-PfH2A). Furthermore, by creating an inducible knock-down system of the endogenous *Pf*SR1, we revealed that *Pf*SR1 is essential for the parasite’s ability to overcome DNA damage as well as for resisting exposure to artemisinin.

## Results

### Stage dependent analysis of PfSR1 interactome points towards its involvement in multiple processes of RNA metabolism and DNA damage response (DDR)

We have previously shown that *Pf*SR1 regulates alternative splicing and RNA levels in *P. falciparum*(4,5). To better understand the mechanisms by which *Pf*SR1 functions as a regulator of gene expression in *P. falciparum*, we were interested in identifying its interacting proteins throughout its intra-erythrocytic development (IDC). We transfected NF54 parasites with an expression vector that allows a fine-tuned over-expression of *Pf*SR1 fused with a Halo-tag at the N-terminus (Fig. S1A). This episomal system allowed us to perform gradual over-expression of *Pf*SR1 by increasing blasticidin concentrations (4), and to perform highly specific pull down assays of *Pf*SR1 interacting proteins *in vivo*(16). As a first step, we gradually over expressed *Pf*SR1-Halo by selecting on increasing blasticidin concentrations (2, 6, & 10μg/ml) and determined the optimal selection pressure needed to detect episomal expression (6μg/ml; Fig. S1B). We then performed stage dependent pull down assays on early and late stage parasites expressing either *Pf*SR1-Halo or a mock plasmid expressing only the Halo-tag (Fig. S1C-D). To identify proteins that were significantly enriched in the fractions recovered from *Pf*SR1-Halo, we performed proteomic analysis on both parasite populations by Liquid Chromatography followed by Mass Spectrometry (LC MS/MS). Proteins that were enriched at least 8 fold in the *Pf*SR1-Halo parasites in 2 out of 3 biological replicates are listed in Table S1. Remarkably, it appears that most of the interactions of *Pf*SR1 with other proteins occur at the early stages of the parasites’ IDC (20 hours post invasion, hpi) and only a few occur at late stages (36 hpi; Fig. 1A). We found that *Pf*SR1 interacts with proteins predicted to function at several levels of RNA metabolism, from transcription and chromatin organization, through splicing and maturation, to translation (Fig. 1B). Furthermore, *Pf*SR1 also interacts with several proteins implicated in DNA damage response (DDR). We further used computational analysis, using the string data base (https://string-db.org/) to predict the protein-protein interaction networks among *Pf*SR1 interacting partners based on known and predicted structure and function of these proteins (Fig. 1C). This analysis, which predicts possible interactions among proteins involved in chromatin organization, transcription and splicing, ribosome biogenesis and DNA damage response, provides an additional indication that *Pf*SR1 may play a role in these processes in addition to its role in splicing and alternative splicing.

**Figure 1:**
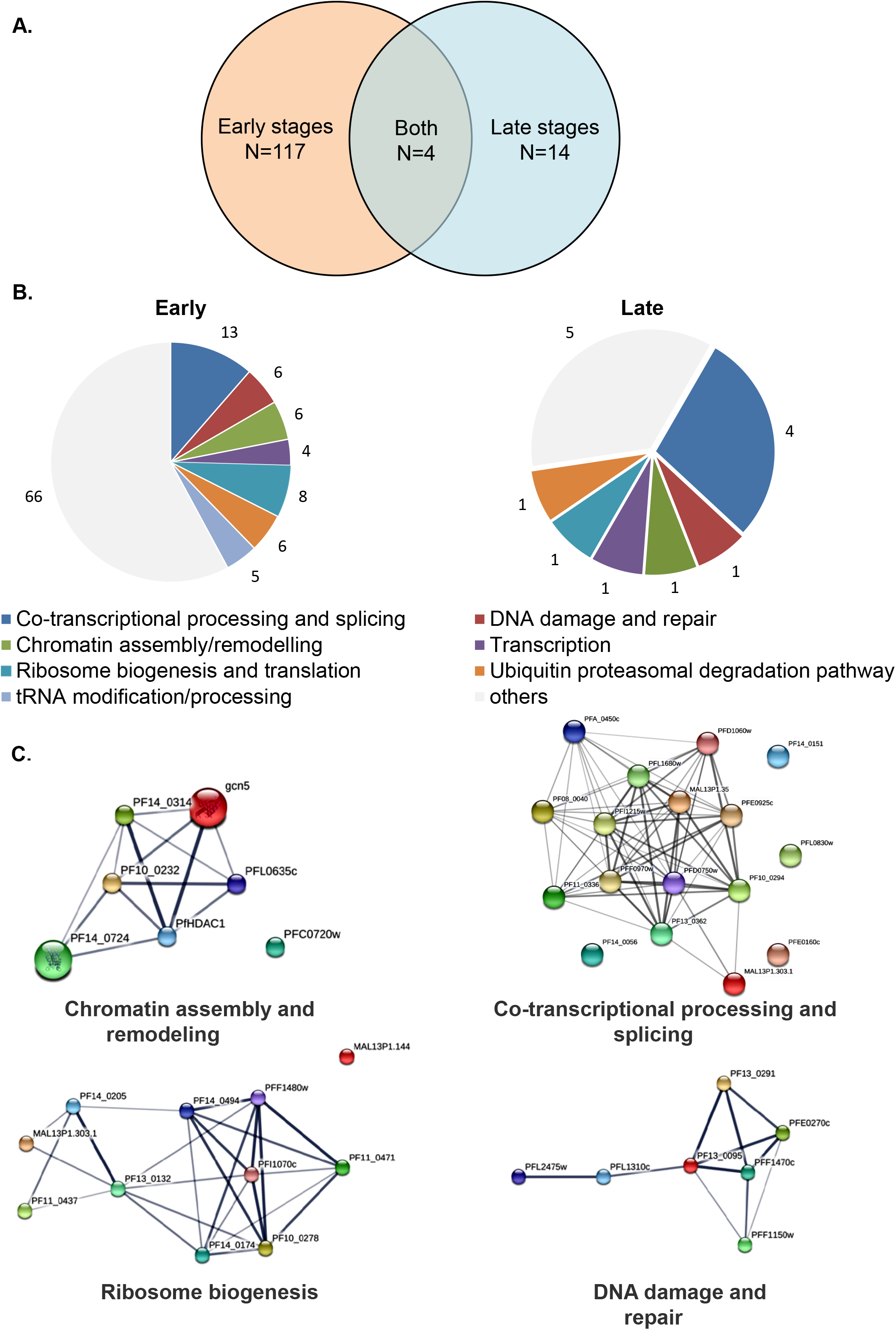
*Pf*SR1 interactome is stage dependent and enriched with proteins involved in several processes of RNA metabolism and DNA damage repair. **(A).** A Venn diagram of stage-dependent *Pf*SR1 interactome showing that most interactions (117 proteins) occur at the first 20h of IDC and only a few (14 proteins) at late stages of IDC **(B).** Pie chart of putative gene annotation of the proteins predicted to be involved in RNA metabolism and DNA damage repair that were specifically enriched in the *Pf*SR1-Halo pull down versus control Halo pull down in early (left) and late stage (right) parasites. The number of genes in each group is presented. Gene annotations were derived from PlasmoDB using annotated GO process. **(C).** Protein-protein interaction networks of *Pf*SR1 interacting proteins. Interaction networks among proteins of different functional groups were calculated using the STRING database. Line thickness indicates the probability of the predicted interactions.

### PfSR1 expression is required for the parasite’s ability to recover from DNA damage

SR proteins were implicated in several processes of RNA metabolism, but very little is known regarding their possible role in DDR. Intrigued by the interactions of *Pf*SR1 with proteins predicted to be involved in DDR (Fig. 1 & Table S1), we decided to investigate its function in these processes. Towards this aim we used the CRISPR / cas9 system to create a transgenic line in which the 3’ terminus of the endogenous open reading frame is replaced and fused with an HA epitope tag and the *glmS* ribozyme (Fig. S2). This parasite line enables to quantify and visualize the endogenous *Pf*SR1 as well as to induce expression knock-down by adding glucosamine (GlcN) to the culture media as described (17). After isolating a clonal population of the transgenic line, we determined that incubation of the parasites with 5mM GlcN for 72h results in an almost complete knock-down of *Pf*SR1 expression (Fig. 2A). Furthermore, we observed that on 5mM GlcN, PfSR1 expression decreases over time and is reversible upon GlcN removal from the culture media (Fig. 2B). Under normal growth conditions, the *Pf*SR1-*glmS* parasite line grew at a similar rate to the NF54 wild type population, however, when *Pf*SR1 was knocked down by GlcN it showed a slight reduction in its growth rate (Fig. 2C & D).

**Figure 2:**
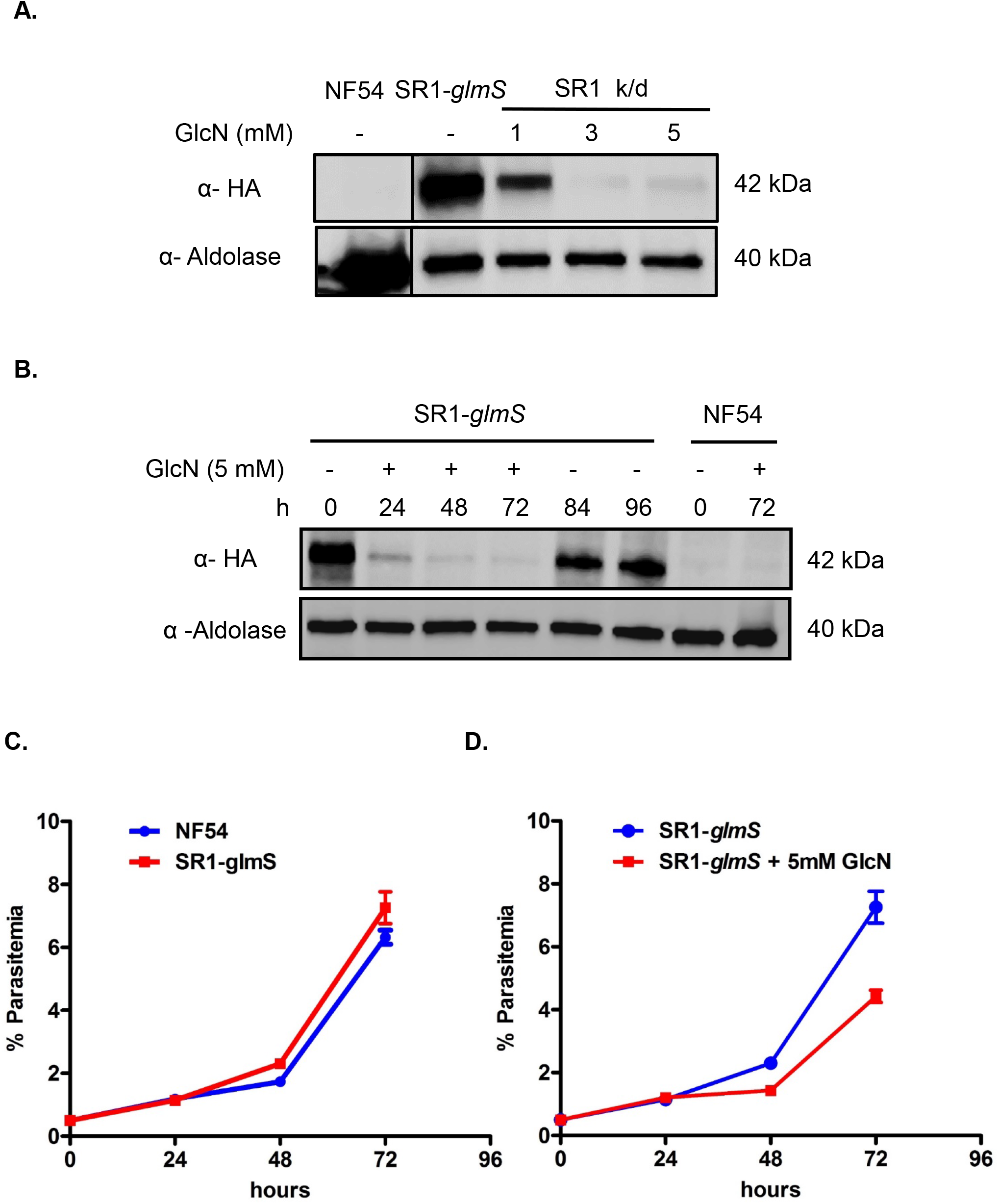
Inducible knock down of endogenous *Pf*SR1. **(A).** Increasing concentration of GlcN for 72h cause decreased expression of endogenous *Pf*SR1 **(B).** Time dependent depletion of *Pf*SR1 in parasites growing on 5 mM GlcN over a 72h time course, which could be reversed by removing GlcN. **(C).** *Pf*SR1-*glmS* and NF54 parasite lines have a similar growth on regular media **(D).** Parasites in which *Pf*SR1 was knocked-down have a delayed growth rate. All experiments were performed in 3 independent biological replicates.

We then used this transgenic line to determine if *Pf*SR1 is involved in DDR. In yeast and mammals, a double strand break (DSB) triggers the DDR that rapidly leads to phosphorylation of the core histone isoform H2A.X to form γ-H2A.X, which marks the site of damaged DNA (18). We have recently demonstrated that in *P. falciparum* that lacks the H2A.X variant, the canonical PfH2A (PF3D7_0617800) is phosphorylated on serine 121 upon exposure to sources of DNA damage and that phosphorylated PfH2A is recruited to foci of damaged chromatin shortly after exposure to sources of damage (19). The ability to specifically detect the dynamics of PfH2A phosphorylation using an anti γ-H2A.X antibody provides a useful marker for studying DNA damage response in *P. falciparum*(19). As a first step, we were interested to confirm that DNA damage could be detected in our transgenic lines similar to what we have previously shown in NF54 parasites. We exposed the *Pf*SR1-*glmS* transgenic line to different doses of X-ray irradiation (1000 & 6000 rad), and found that the levels of γ-PfH2A gradually increased following parasites’ irradiation), however, the levels of *Pf*SR1 expression were not significantly elevated (Fig. 3A). To further demonstrate direct measures of DNA damage in response to increased levels of X-ray irradiation we performed a TUNEL assay and visualized DNA fragmentation in the parasites’ nuclei. We clearly show that in untreated parasites less than 10 percent of the parasites had signal, however, in parasites exposed to 1000 and 6000 rad, a strong signal was observed in over 50 and 90 percent of the nuclei, respectively (Fig. 3B). These data demonstrate that X-ray irradiation induces DNA damage in non-replicating ring stage *Pf*SR1-*glmS* parasites and that γ-PfH2A levels are elevated during DDR. Interestingly, in both *Pf*SR1-*glmS* and NF54 parasites, we observed an increase in γ-PfH2A levels 15 minutes following X-ray irradiation (6000 rad), irrespective of GlcN treatment (Fig. 3C). We then tested if downregulation of *Pf*SR1 expression is associated with DDR *in vivo* over a longer period of time and found that the levels of γ-PfH2A are elevated approximately one week following PfSR1 knock-down (Fig. 3D), while only basal levels of γ-PfH2A is observed in NF54 parasites (Fig. S3).

**Figure 3:**
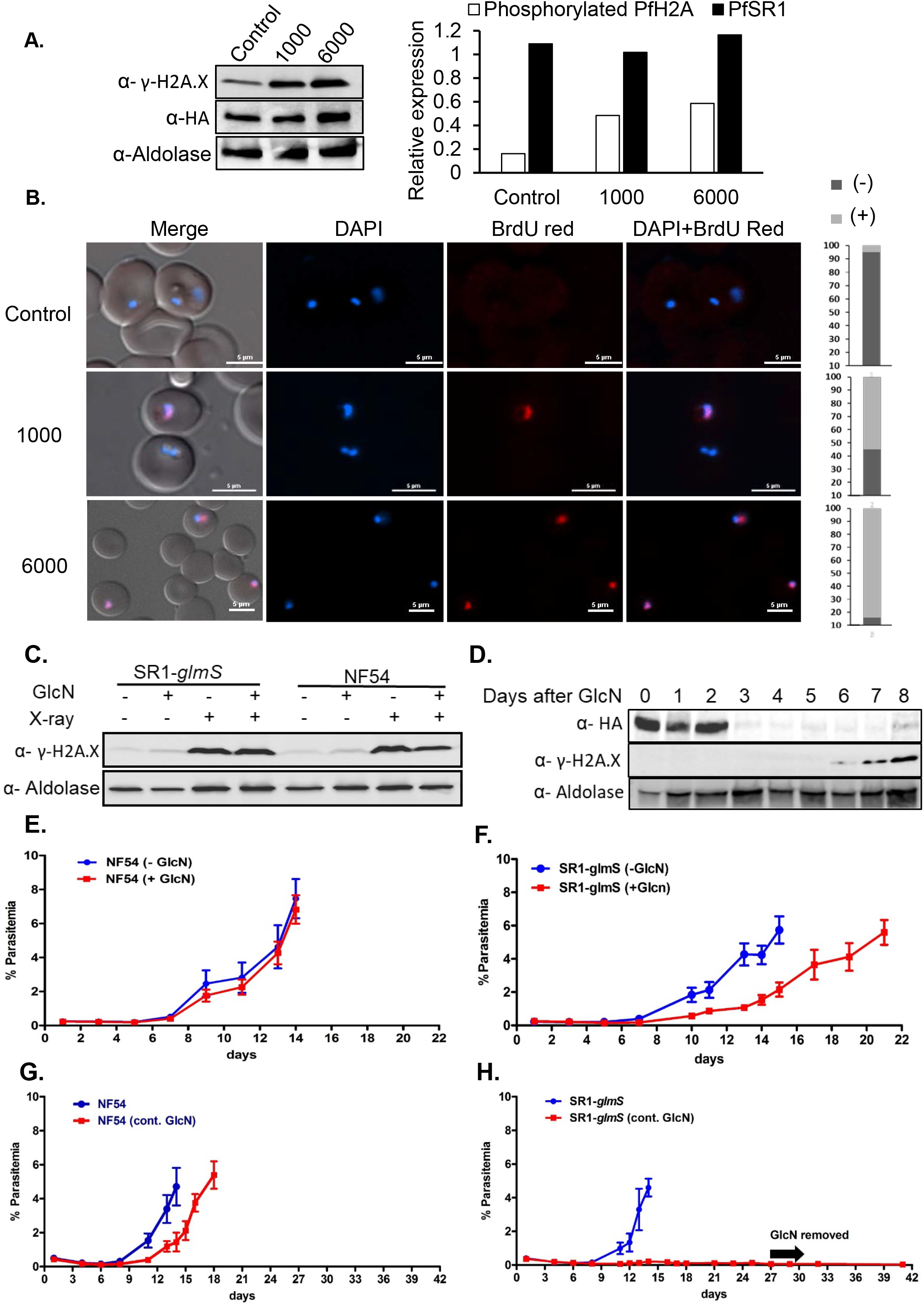
*Pf*SR1 is essential for parasite recovery from DNA damage cause by X-ray irradiation. **(A).** Exposure of *Pf*SR1-*glmS* parasite lines to increasing levels of X-ray irradiation is associated with increased levels of γ-PfH2A. **(B).** DNA fragmentation imaging by TUNEL assay of parasites exposed to increasing levels of X-ray irradiation demonstrating increased levels of DNA breaks. Quantification of the percentage of TUNEL positive and negative nuclei in each treatment is presented on the right (n=100). Scale bar is 5 μm. **(C).** Western blot analysis demonstrating that the levels of γ-PfH2A are elevated 15 minutes after X-ray irradiation regardless of GlcN treatment or parasite line (5mM for 72 h prior to irradiation). **(D).** Western blot analysis of *Pf*SR1-*glmS* parasite line growing on GlcN over time, indicating that γ-PfH2A accumulates approximately one week after *Pf*SR1 down-regulation. **(E).** Growth curves of wild type NF54 parasites exposed to a near lethal dose of X-ray irradiation of 6000 rad. Parasites were grown either on regular media (-GlcN) or with media added with 5mM GlcN for 72h and washed immediately before irradiation (+ GlcN). **(F).** Growth curves of *Pf*SR1-*glmS* parasites exposed to 6000 rad X-ray irradiation and grown either on regular media (-GlcN) or with media supplemented with 5mM GlcN for 72h and washed immediately before irradiation (+ GlcN). **(G).** Growth curves of wild type NF54 parasites exposed to 6000 rad X-ray irradiation and grown either on media supplemented continuously with 5mM GlcN (cont. GlcN) or on regular media. **(H).** Growth curves of *Pf*SR1-*glmS* parasites exposed to 6000 rad X-ray irradiation and grown either in media supplemented continuously with 5mM GlcN (cont. GlcN) or in regular media. Each of the curves represents the average parasitemia of 3 biological replicates at each timepoint. Error bars represent standard errors.

To determine if *Pf*SR1 is essential for the parasite’s ability to recover from DNA damage, we irradiated parasites with lethal (6000 rad) doses and followed their rate of recovery as previously described (20). We found that NF54 parasites recover approximately 13 days after X– ray irradiation, and that pre-incubation with GlcN for 72h prior to irradiation did not affect their growth rate (Fig. 3E). However, in the *Pf*SR1-*glmS* line, a 5-6 day delay in parasite recovery was observed following *Pf*SR1 knocked-down for 72h prior to irradiation as compared to those which were not pre-incubated with GlcN (Fig. 3F). In this set of experiments, GlcN was removed from the culture media just before irradiation, thus allowing the parasites (in which *Pf*SR1 was knockdown) to recover from the damage caused by X-ray irradiation while expressing *Pf*SR1. To determine if *Pf*SR1 is indeed essential for the recovery of parasites from X-ray irradiation we performed similar experiments, only this time we compared the recovery of parasites in the presence of GlcN over the entire course of the experiment (i.e. under continuous knock-down of *Pf*SR1) even after X-ray irradiation. Strikingly, while NF54 parasites recovered on continuous growth with GlcN (Fig. 3G), *Pf*SR1-*glmS* parasites could not recover from the damage caused by the lethal dose of irradiation unless *Pf*SR1 was expressed (Fig. 3H). To exclude the possibility that the observed phenotype is due to accumulated growth defects of long term *Pf*SR1-depletion, we show that PfSR1-depleted line maintains their delayed growth rate constantly over a long period of time without irradiation (Fig. S4). Altogether, these data indicate that *Pf*SR1 expression is essential for *P. falciparum* parasites to overcome the damage caused by X-ray irradiation and point towards its involvement in the DNA damage response.

### PfSR1 is recruited to the sites of DNA damage

The fact the γ-PfH2A is recruited to damaged chromatin shortly after exposure to sources of DNA damage (19) enabled us to visualize a possible association between *Pf*SR1 and the nuclear site of DNA damage. We used the transgenic *Pf*SR1-*glmS* line to visualize the HA-tagged endogenous *Pf*SR1 and determine if it co-localizes with γ-PfH2A. We found a strong association (78/83 independently counted nuclei) between *Pf*SR1 and the site of DNA damage in both early and late stages parasites (Fig. 4A). In addition, we found that in irradiated ring stage parasites *Pf*SR1 co-localizes with *Pf*Rad51 (Fig. 4B, 39/50 independently counted nuclei), providing additional support for its recruitment to the site of DNA damage. Furthermore, we performed histone extraction from irradiated parasites, and found that *Pf*SR1 is associated with histones (Fig. 4C). Encouraged by this observation, we were interested to determine whether *Pf*SR1 interacts with the phosphorylated PfH2A at the damaged loci. To this end, we used the anti γ-H2A.X antibody to perform co-immuno precipitation (Co-IP) on nuclear extracts of irradiated parasites. Western blot analysis of this Co-IP experiment shows that both γ-PfH2A and *Pf*SR1 were present in the IP eluted fraction (Fig. 4D), indicating that they were co-immuno precipitated. Moreover, as expected, only a small amount (compared with the input) of the core histone H3 was also pulled down, and the cytoplasmic aldolase could not be detected. These data indicate that *Pf*SR1 is recruited to the site of DNA damage where it interacts with γ-PfH2A.

**Figure 4:**
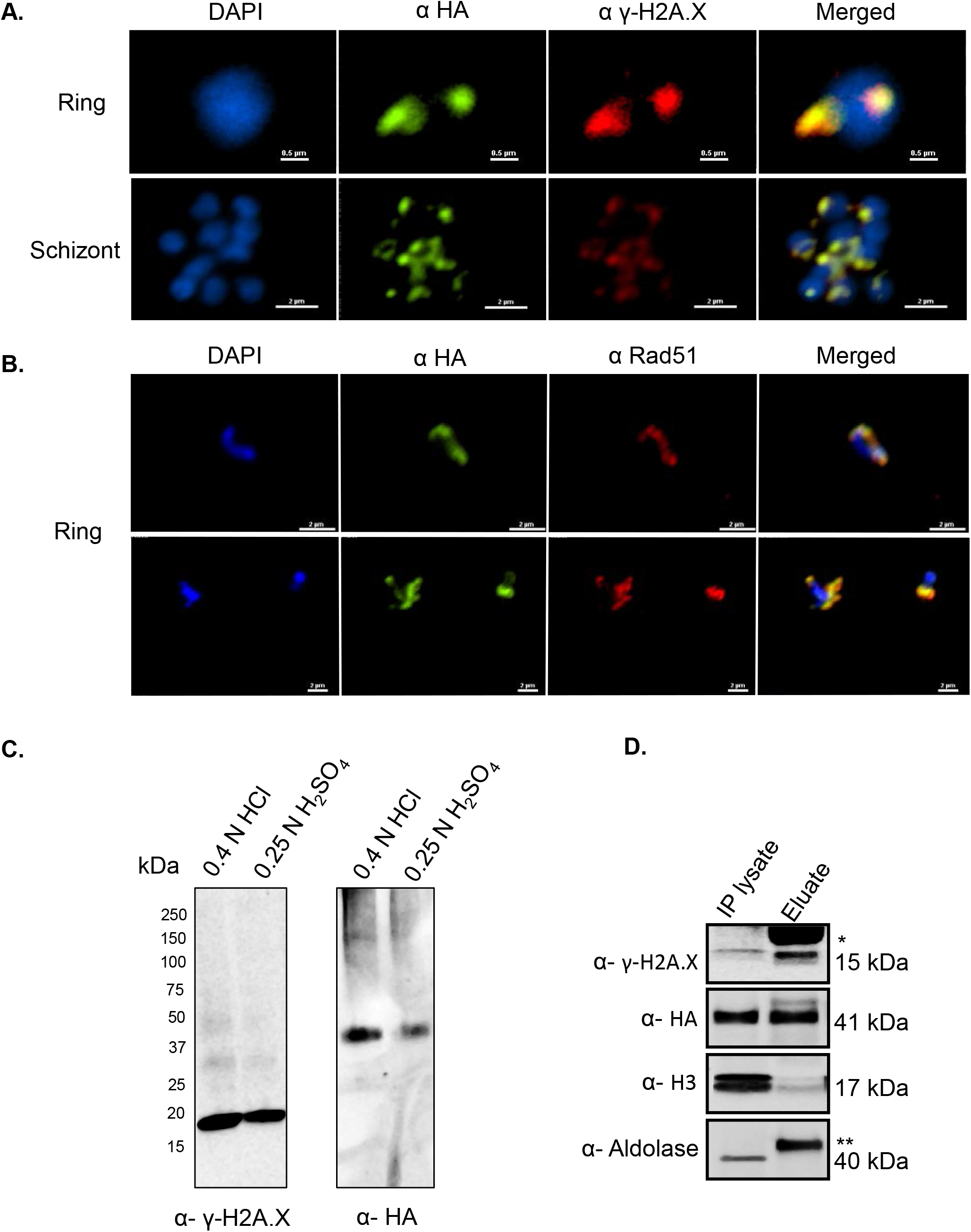
*Pf*SR1 is recruited to the site of damaged DNA and interacts with phosphorylated H2A. **(A).** Immunofluorescence assay demonstrating that *Pf*SR1 is associated with γ-PfH2A in the nucleus of early (upper panel) and late stage (lower panel) *Pf*SR1-*glmS* parasites 15 minutes after X-ray irradiation. **(B).** Immunofluorescence assay demonstrating that *Pf*SR1 co-localize with *Pf*Rad51 in the nucleus of ring stage *Pf*SR1-*glmS* parasites 15 minutes after X-ray irradiation. **(C).** Total histone extraction associate *Pf*SR1 with histones. Histone extraction from irradiated (6000 rad) PfSR1-*glmS* parasites was performed as described (19). Western blot analysis using anti γ-H2A.X antibody (left) and anti HA (right) detected both *Pf*SR1 and γ-PfH2A in histone fractions extracted with either HCl or H2SO4. **(D).** *Pf*SR1 interacts with γ-PfH2A as detected by Co-IP experiment using anti γ-H2A.X antibody. Western blot analysis for the presence of γ-PfH2A, *Pf*SR1 (α HA), core histone H3 (α H3), and cytosolic aldolase in the input (IP lysate) and the pulled down fraction (eluate) demonstrate that *Pf*SR1 was pulled down with the anti γ-H2A.X antibodies. Light chain of primary antibody is marked with * and heavy chain is marked with **.

### PfSR1 is essential for repairing DNA damage

The association of *Pf*SR1 with damaged chromatin and its essentiality for the ability of parasites to recover from exposure to X-ray irradiation led us to test its involvement in DNA damage repair. We have recently shown that PfH2A phosphorylation is dynamic and over time, as the parasite activates the repair machinery, this phosphorylation is removed. Thus, these phosphorylation dynamics were used to establish a direct DNA repair assay in *P. falciparum* (19). To exclude the possible effect of GlcN on the parasite’s ability to activate the DNA repair machinery, NF54 parasites were exposed to sub-lethal levels of X-ray irradiation (1000 rad), put back in culture to allow the parasites to repair the damaged DNA (Fig. 5A). We observed that in NF54 parasites growing in either regular media (left) or in media added with 5 mM GlcN (right), the levels of γ-PfH2A increased 15 minutes after irradiation. However, after 3 and 6 hours the levels of γ-PfH2A decreased back to the basal level of parasites that were not irradiated, indicating that the DNA repair process was initiated (Fig. 5A). We then performed the same assay on the *Pf*SR1-*glms* transgenic line. We found that in the *Pf*SR1-*glms* parasites growing in regular media and expressing *Pf*SR1 (Fig. 5B, left panel) there is an increase in γ-PfH2A levels 15 minutes after irradiation and a decrease 3 and 6 hours later indicating that during that time the parasites were able to activate the DNA repair machinery. In marked contrast, PfSR1-*glms* parasites growing in the presence of 5 mM GlcN where PfSR1 expression is downregulated, the levels of γ-PfH2A remained constant, similar to the levels measured immediately after irradiation (Fig 5B, right panel). This data indicates that in the absence of *Pf*SR1 expression, *P. falciparum* parasites are impaired in their ability to repair DNA damage.

**Figure 5:**
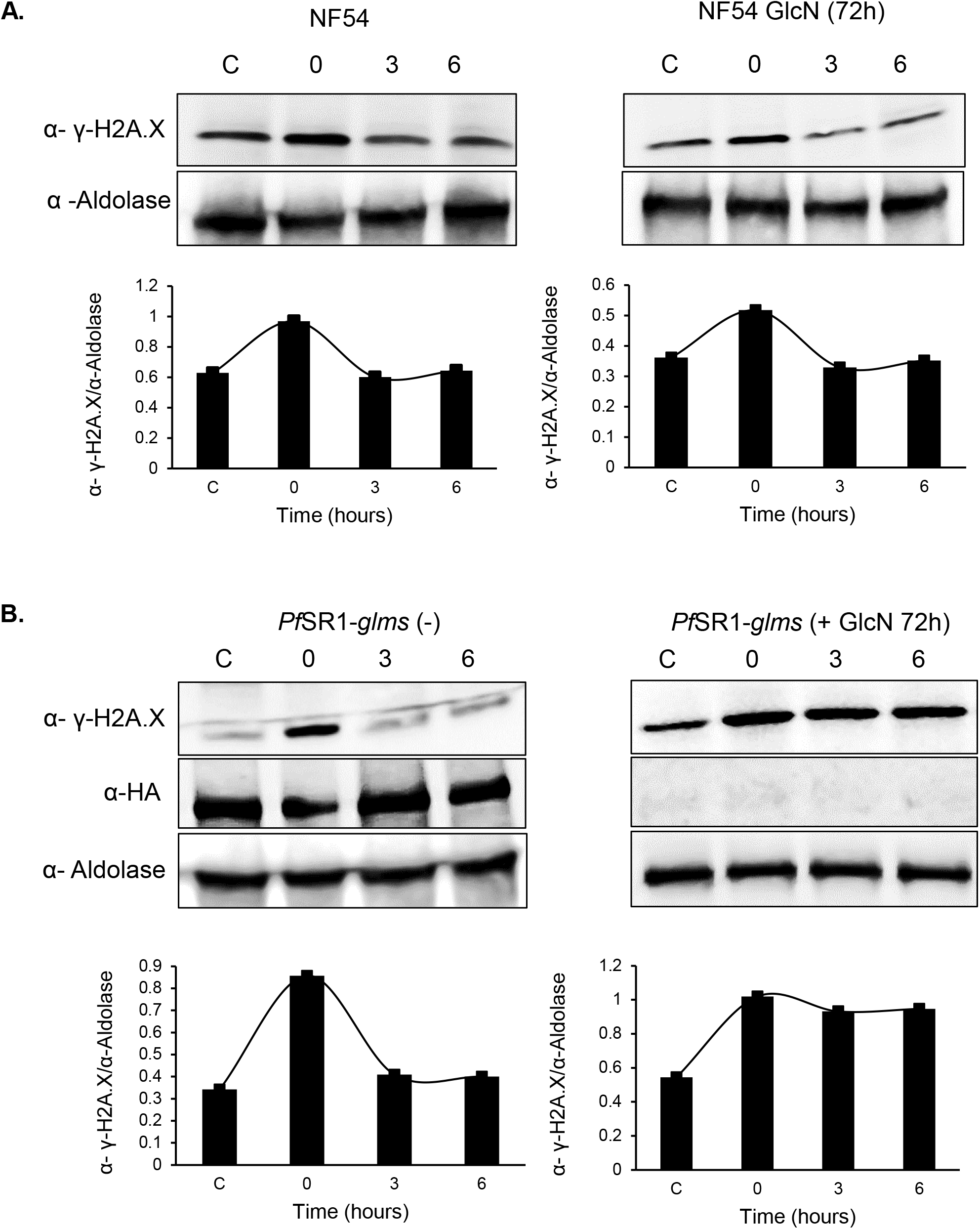
*Pf*SR1 is essential for repairing damaged DNA. **(A).** Upper panels: Western blot analysis of protein extracted 15 minutes, 3 hours and 6 hours (time 0, 3 & 6 respectively) following exposure to a sub lethal dose of X-ray irradiation (1000 rad). Protein was extracted from tightly synchronized ring stage NF54 parasites growing either on regular media (left) or on media supplemented with 5 mM GlcN (right). Lower panels: Densitometry quantification of the ratio between the Western blot signals obtained for γ-PfH2A and aldolase. These analyses demonstrate that the level of γ-PfH2A, which increases 15 minutes after irradiation, is reduced 3 hours later regardless of GlcN. **(B).** Upper panels: Western blot analysis of protein extracted 15 minutes, 3 hours and 6 hours (time 0, 3 & 6 respectively) following exposure to a sub lethal dose of X-ray irradiation (1000 rad). Protein was extracted from tightly synchronized ring stage *Pf*SR1-*glmS* parasites growing either on regular media (left) or on media supplemented with 5 mM GlcN (right). Lower panels: Densitometry quantification of the ratio between the Western blot signals obtained for γ-PfH2A and aldolase. These analyses demonstrate that in the *Pf*SR1-*glmS* parasites DNA damage could only be repaired in parasite growing on regular media and not those growing on GlcN in which *Pf*SR1 is down regulated.

### PfSR1 is essential for the parasite’s ability to recover from artemisinin exposure

The cytotoxic effect of artemisinin, the first line antimalarial chemotherapeutic, was recently attributed to its ability to cause DNA damage mediated by reactive oxygen species (ROS) (21). Moreover, short exposure of parasites to artemisinin elicits a DNA damage response similar to that induced by the alkylating agent, methyl methanesulphonate (MMS), which further supports artemisinin activity as a DNA damaging agent (9). We found that the levels of γ-PfH2A are elevated following exposure of tightly synchronized ring stage parasites to artesunate, the semisynthetic derivative of artemisinin (Fig. 6A), and that already 15 minutes after artesunate treatment, DNA fragmentation could be visualized by TUNEL in more than 95% of the parasites (Fig. 6B). Considering the critical role that *Pf*SR1 plays in *P. falciparum* DDR, we tested if it is required for parasite recovery from artemisinin pressure. To this end, we used the *Pf*SR1-*glmS* parasites and compared their ability to resist treatment with artesunate (ART), when the endogenous *Pf*SR1 was expressed or when it was knocked-down by GlcN incubation 72h prior to artesunate treatment. Tightly synchronized ring stage parasites were incubated with 700 nM artesunate for 6 hours, then washed and put back into culture on regular media as previously described (22). To evaluate the recovery rate of each parasite population from artesunate exposure, the level of parasitemia was measured daily by flow cytometry. We found that when *Pf*SR1 was expressed, parasites recovered approximately 18 days after artesunate treatment, while the recovery of parasites in which *Pf*SR1 was knocked-down prior to artesunate treatment was delayed (Fig. 6C). Strikingly, parasites in which *Pf*SR1 was continuously downregulated by the addition of GlcN to the media were unable to recover from artesunate treatment, and no parasites were detected even after 21 days. These results imply that *Pf*SR1 expression is essential for parasite recovery from artesunate treatment.

**Figure 6:**
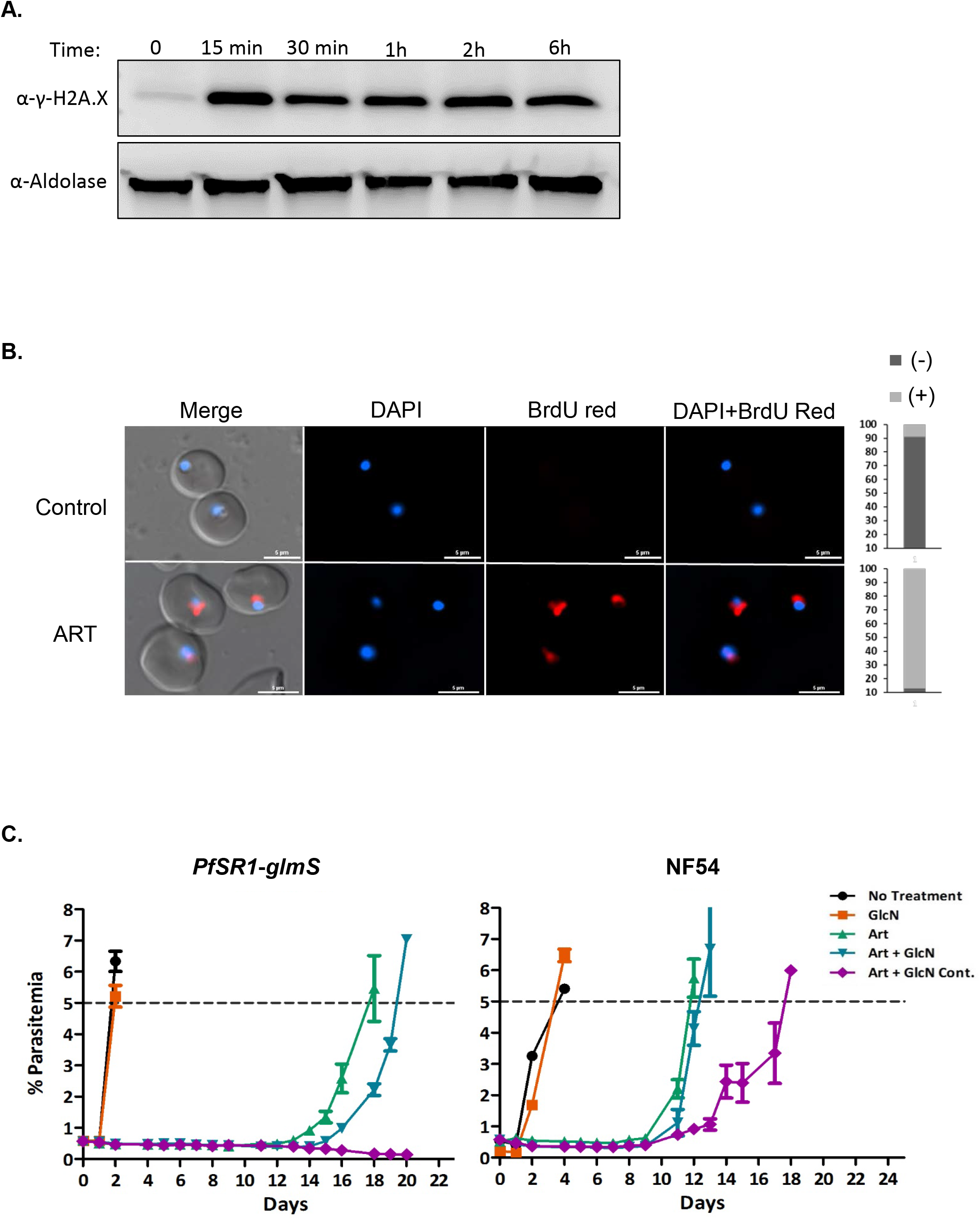
*Pf*SR1 expression is required to overcome artesunate exposure. **(A).** Western blot analysis demonstrating the increase in γ-PfH2A levels following exposure of tightly synchronized ring stage parasites to 700 nM artesunate for different time periods. **(B).** DNA fragmentation imaging by TUNEL assay of parasites exposed to 700 nM artesunate for 15 minutes, demonstrating increased levels of DNA breaks. Quantification of the ratio of TUNEL positive and negative nuclei in each treatment is presented on the right (n=100). Scale bar is 5 μm. **(C).** Growth curves of tightly synchronized ring stage *Pf*SR1-*glmS* (left) and NF54 parasites (right), treated with 700 nM artesunate for 6h. GlcN was either not added (Art, 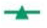), added 72h prior to artesunate treatment (Art+GlcN, 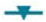), or added continuously throughout the entire recovery period (Art+GlcN cont. 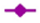). Parasites growing on regular media without any treatment (no treatment, 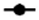) and parasites treated only with GlcN (GlcN, 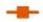) are presented as controls. The results presented are the average parasitemia of 3 biological replicates measured daily. Error bar represents standard errors.

## Discussion

All eukaryotic organisms must maintain their genomic integrity and are continuously exposed to challenges threatening to damage their DNA (7). *Plasmodium* parasites that replicate by schizogony within human red blood cells are particularly prone to the most common sources causing DNA damage, such as replication defects and oxidative stress. In recent years, RNA binding proteins (RBPs) were implicated as important keepers of genome stability and as active players in the DDR (6,7,23). Among those, several splicing factors, which in addition to their canonical role in splicing regulation, were shown to play a direct role in preventing DNA damage during transcription and cellular segregation, as well as to be actively involved in DNA repair (7,24). In the current study, we found that the interactome of *Pf*SR1, an alternative splicing factor of *Plasmodium falciparum*, includes putative proteins that could be implicated in DDR. These findings led us to hypothesize that in addition to its role in regulating splicing and RNA metabolism, *Pf*SR1 is also involved in DDR. Using endogenously tagged regulatable *Pf*SR1, we have demonstrated that *Pf*SR1 is recruited to foci of damaged DNA, where it interacts with γ-PfH2A, a marker for damaged DNA. Furthermore, we showed that *Pf*SR1 is essential for the parasite’s ability to activate the DNA damage repair mechanism and overcome induced DNA damage as well as exposure to artemisinin, the first line antimalarial drug.

At different stages of their life cycle, *Plasmodium* parasites replicate their haploid genome multiple times through consecutive mitosis cycles called schizogony, which makes them particularly prone to error during DNA replication. In addition, blood stage parasites live in a highly oxygenated environment, and while digesting large amounts of hemoglobin in their food vacuole they produce heme and hydroxyl radicals which are potent DNA damaging agents (25). Therefore, it is reasonable that the parasite has evolved efficient mechanisms to protect its genome integrity, which will allow it to proliferate in such conditions. Although little is known on DDR in *Plasmodium* parasites, their genome contains orthologues to many of the proteins involved in homologous recombination (HR), microhomology-mediated end joining (MMEJ) (26) and mismatch repair machinery (12). Surprisingly, it appears that none of the components of the canonical non-homologous end joining (C-NHEJ) pathway could be identified in these parasites. Recently, Kirkman et al, created a transgenic line containing a system for inducing targeted double strand breaks (DSBs) in *P. falciparum*. They showed that malaria parasites utilize both HR and an alternative end joining pathway to maintain genome integrity, with preference to the alternative end-joining pathway as a primary method for repairing DSBs in the absence of heteroallelic homologous sequences that can serve as templates for HR (10).

In addition to the intrinsic metabolic oxidative stress sources, *Plasmodium* parasites are exposed to oxidative substances released from immune cells as a response to the infection, as well as to reactive oxygen species mediated by chemotherapeutic agents used to treat malaria which can significantly disrupt genomic integrity (21,27,28). In this regard, activated artemisinin and its derivatives react promiscuously with nucleophile-harboring cellular components, leading to alkylation of DNA and proteins which may cause protein misfolding and DNA damage (29). Indeed, the recent association of the K13 mutation with artemisinin resistance was also accompanied by evidence for enhanced unfolded protein response (UPR) in the resistant mutant lines (30). In addition, artesunate was shown to induce oxidative DNA damage and response in mammalian cancer cells (31–33) as well as DNA damage in malaria parasites, which is mediated by reactive oxygen species (21). Recently, it was demonstrated that exposure of *P. falciparum* to artemisinin induced DDR, involving both transcriptional and epigenetic changes that are similar to the parasite response to the DNA damaging agent methyl methanesulphonate (MMS) (9). It is therefore likely that the cytotoxicity of artemisinin to malaria parasites involves direct DNA damage. The novel function of *Pf*SR1 in protecting the parasites from DNA damage and its critical role in enabling the parasite to overcome exposure to artemisinin, supports reported evidence that artemisinin and its derivatives damage genome integrity of the *Plasmodium* parasites.

DNA damage was shown to affect expression, localization and post-translational modifications of many splicing factors which in turn are involved in regulation of the DDR at different levels (6,23). For example phosphorylation of SRSF1, the human orthologue of *Pf*SR1, is regulated during the DDR, shifting the alternative splicing pattern of target genes to control cell survival (34). However, to the best of our knowledge, we demonstrate for the first time that *Pf*SR1 is recruited to the site of damaged chromatin where it interacts with γ-PfH2A. This association may suggest that *Pf*SR1 plays a functional role at the site of damaged DNA in preserving genome integrity after exposure to irradiation or artesunate. This conclusion does not exclude the possibility that *Pf*SR1 is also involved in other indirect regulatory pathways in the DDR machinery through regulation of gene expression.

In addition to proteins involved in DDR, the interactome of *Pf*SR1 contains many additional proteins involved in several processes of RNA metabolism ranging from chromatin remodeling, transcription, splicing and translation. Previously, we provided evidence that in addition to its role as an AS factor, *Pf*SR1 binds to specific RNA motifs and regulates the steady state RNA levels of a subset of RNA molecules (5). Moreover, *Pf*SR1 localizes mainly to the nucleus at the early stages of the IDC, while in late stages it is located primarily in the parasites’ cytoplasm (4). This dynamic localization indicates that *Pf*SR1 may have multi-faceted biological functions both in the nucleus and the cytoplasm. In higher eukaryotes, SR proteins are recruited to transcriptionally active chromatin by the C-terminal domain of RNA Pol II during transcription elongation, and the initial steps of RNA processing such as pre-mRNA splicing are coupled with transcription (35,36). Coupling between transcription and mRNA processing depends on chromatin structure and its accessibility to splicing regulators (37). In agreement with these findings, we found that *Pf*SR1 is associated with Pol II as well as with transcription elongation factors and additional chromatin regulators, suggesting that transcription and splicing could be coupled in *P. falciparum* as well. In addition, direct and functional interactions of the SR proteins with specific chromatin modifications, were shown to be essential for proper cell cycle progression (38). Our data show that downregulation of *Pf*SR1 caused a delay in parasite growth rate, however, further investigation is required to determine if *Pf*SR1 interactions with chromatin play a role in the cell cycle progression of malaria parasites. Besides its interaction with components of the transcriptional machinery, *Pf*SR1 also interacts with another SR protein (PF3D7_0503300), which shares significant homology with the mammalian SRSF2 and SRSF12. Interestingly, SRSF2 was shown to be restricted to the nucleus where it associates with DNA (39) and recently was shown to play a role in regulating transcriptional activation by releasing paused Pol II on gene promoters (40). It will be interesting to further investigate the possibility that *Pf*SR1 is involved in transcriptional regulation.

SR proteins were also shown to be involved in mRNA export as well as in regulation of cytoplasmic processes, such as mRNA decay and translation. In the late stages of the IDC, *Pf*SR1 was found to be localized mainly to the cytoplasm (4), but its cytoplasmic function was never investigated. Interestingly, we found that most of the *Pf*SR1 interactions with other proteins occur at 20-24 hours post infection. During this first phase of the IDC, *Pf*SR1 interacts with proteins involved in ribosome biogenesis and translation. This could indicate that *Pf*SR1 shuttling to the cytoplasm begins already at that phase, and that *Pf*SR1 might also be involved in the translation process. Cytoplasmic SR proteins were found to interact with actively translating ribosomes and are capable of stimulating protein synthesis by promoting mRNA entrance to polysomes (41,42), stimulating mTOR, and/or enhancing phosphorylation of 4E-BP1 (43,44). The interactions of *Pf*SR1 with polyadenylated RNA binding protein, the translation initiation factor EIF-2B and other ribosomal proteins may indicate that similar to SRSF1, it could contribute to translational enhancement in the cytoplasm. Interestingly, 36% of the binding partners of SRSF1 are ribosomal proteins, and its specific interaction with RPL5 induces cellular senescence (45), which is essential for DNA damage repair. Altogether, our data point towards a novel role of *Pf*SR1 in protecting *P. falciparum* parasites from different sources of DNA damage and opens new avenues for investigation into the mechanisms involved.

## Materials and Methods

### Parasite culture and parasitemia counts

All parasites used were derivatives of the NF54 parasite line and were cultivated at 5% hematocrit in RPMI 1640 medium, 0.5% Albumax II (Invitrogen), 0.25% sodium bicarbonate, and 0.1 mg/ml gentamicin. Parasites were incubated at 37°C in an atmosphere of 5% oxygen, 5% carbon dioxide and 90% nitrogen. Parasite cultures were synchronized using percoll/sorbitol gradient centrifugation as previously described (46,47). Briefly, infected RBCs were layered on a step gradient of 40%/70% percoll containing 6% sorbitol. The gradients were then centrifuged at 12,000g for 20 minutes at room temperature. Highly synchronized, late stage parasites were recovered from the 40%/70% interphase, washed twice with complete culture media and placed back in culture. The level of parasitemia was calculated by Flow Cytometry. For flow cytometry, aliquots of 50μl parasite cultures were washed in PBS and incubated 30 min with 1:1000 SYBR Green I DNA stain (Life Technologies). The fluorescence profiles of infected erythrocytes were measured on CytoFLEX (Beckman Coulter) and analyzed by the CytExpert software.

### Plasmid construction

In order to express PfSR1 fused with a Halo-tag, a Halo-tag sequence was amplified using the primers nHalo-F-nHaloR (GGCGACTAGTCCATGGCAGAAATCG - CCGCGAGCTCTGAATTCGGAAGCGATC) and cloned into the expression vector pHBIRH (4) using SpeI – SacI (essentially replacing renilla luciferase and the hrp2 3’ and introducing AsiSI restriction site) to generate the pHBInHalo vector. Then *pfsr1* (PF3D7_0517300) was amplified together with 1000 bp of its 3’ UTR using the primers n*Pf*SR1F – n*Pf*SR1-UTR-R (5’-CCAGCGATCGCAATGGTTATACGTGAAAGT-3’ – 5’-CCGGAGCTCATATCTATATTTTTGTAAA-3’) and cloned into pHBInHalo using *AsisI–SacI* to generate pHBInHaloSR1UTR. To create *Pf*SR1-*glmS* transgenic line we used a modified CRISPR/ cas9 previously described (22). The cas9 nuclease was expressed with pUF1-Cas9. The pL6-*Pf*SR1-HA-*glmS* plasmid, was constructed by amplifying *Pf*SR1 homology arm from the 5’ of its ORF using primers: 5’–AGGTGAATGTGGTCATGCAG-3’ and 5’-ATGTCTTCTTTTATGGGACGATGATG-3’. *Pf*SR1 endogenous 3’ UTR was amplified using the primers: 5’-CGTCCCATAAAAGAAGACATTAGA-3’ and 5’-ATGCTTAAG CAGTGCGAGGCTCTATTATGTG-3’, both fragments were cloned into pL7-PfSAC1-3HA-*glmS*-DHFR using (48) using Not1/Sma1 and Nhel/Afll1 respectively to generate the pL6-*Pf*SR1-3HA-*glmS*-DHFR. For cloning the sgRNA we used the In-Fusion HD Cloning Kit (clontech), with the 20-bp guide RNA (CCAACTCAAGATCATCATCA) surrounded by the 15 bp necessary for In Fusion cloning. The final pL6-*Pf*SR1-3HA-*glmS*-DHFR constructs were made by replacement of the BtgZI-adaptor with the guide RNA sequence. All PCR amplifications were done with high-fidelity Q5 Taq polymerase (New England Biolab).

### Parasite transfection and selection

Parasites were transfected as described (49). Briefly, 0.2 cm electroporation cuvettes were loaded with 0.175ml of erythrocytes and 50 μg of plasmid DNA in an incomplete cytomix solution. Stable transfectants carrying plasmids with an hDHFR-selectable marker were selected on 4 nM WR99210 and those carrying yDHODH were selected on 1.5 μM DSM1. Stable transfectants carrying plasmids with BSD-selectable marker were initially selected on 2μg/mL blasticidin-S (Invitrogen). In order to obtain parasites carrying large plasmid copy numbers, these cultures were then subjected to elevated concentrations of 6-10 μg/ml blasticidin-S, depending on experimental design.

### HaloLinK pull down assay

Halo pull down assay was performed on extracts made from 400-500 ml of tightly synchronized parasite cultures of early (20 hpi) and late (36 hpi) stages. iRBCs were saponin lysed, washed twice in PBS and the parasite pellets were stored at −80°C for at least 30 min. Parasite pellets were then thawed in 200 μL lysis buffer (50 mM Tris-HCl (pH 7.5), 150 mM NaCl, 1% Triton X-100, 0.1% Na deoxycholate) with protease inhibitors cocktail (Promega, cat # G6521), mixed well by pipetting and incubated on ice for 15-30 min. Pellets were then centrifuged at 14,000×g for 5 min at 4°C and the supernatant was transferred into a fresh new tube added with TBS to the final volume of 1 ml (80 μl of this starting material (SM) was kept for Western blotting). Pull down of Halo-*Pf*SR1 interacting proteins was performed using the HaloTag^®^ Protein Purification System (Promega, cat # G1913) according to the manufacturer guidelines. Briefly, HaloLink resin was washed 3-5 times with 800 μl wash buffer (90 μl of 10% NP40 in 18 ml TBS) and the sample was added to the HaloLink resin and incubated in 4°C overnight while rotating. The following day the sample was centrifuged and the resin was washed 3 times in wash buffer while the flow through (FT) was kept for quality control by western blot. To recover the purified proteins of interest (without the resin bound tag), elution was performed by resuspending the resin in 50 μl SDS solution buffer and incubating for 30 min at 1400 rpm at 55°C. The starting materials, flow through fractions and eluates were all checked by Western blot for the presence of the Halo-tag. To verify specific enrichment in the Halo-*Pf*SR1 pull down, the purified proteins were loaded on an SDS-PAGE and silver stained for protein visualization (Pierce Silver Stain Kit, Thermo Scientific). Coomassie stained SDS-PAGE samples were sent for mass spectrometry analysis.

### LC-MS/MS Analysis and Database analysis

The proteins obtained by HaloLink pull down were separated on a 4–20% acrylamide gel (BioRad) and stained with NOVEX Colloidal Blue Staining Kit (Invitrogen). The samples were digested by trypsin, analyzed by LC-MS/MS on Q Exactive plus (Thermo) and identified by Discoverer software version 1.4 against the *Plasmodium* NCBI-NR database and against decoy databases in order to determine the false discovery rate (FDR) using the sequest and mascot search engines. Semi quantitation was done by calculating the peak area of each peptide. The area of the protein is the average of the three most intense peptides from each protein. The results were filtered for proteins identified with at least 2 peptides with 1% FDR. We considered specific *Pf*SR1 enrichment as proteins that were enriched, either more than 8 fold or found only in the Halo-*Pf*SR1 pull down fraction and not in proteins extracted from parasites transfected with the mock plasmid, in at least two replicates.

### Southern blot

Analysis of the integrated construct was performed using Southern blots and diagnostic PCR crossing the integration sites (using primers: P1F 5’-ACGTGAAAGTGTATCGAGAA-3’, P2R 5’-AGTGCGAGGCTCTATTATGTG-3’, and P3R 5’-ATGCCTTTCTCCTCCTGGAC-3’ followed by sequencing. Southern blots were performed according to established protocols (50,51). Briefly, genomic DNA isolated from recombinant parasites was digested to completion by the restriction enzymes *SpeI* and *NheI* and subjected to gel electrophoresis using 1% agarose gel in Tris / Borate / EDTA (TBE). The DNA was transferred to a high-bond nitrocellulose membrane by capillary action after alkaline denaturation. DNA detection was performed using DIG High Prime DNA Labeling and Detection starter kit (Roche). The *glmS* sequence amplified from pL6-HA-*glmS* using 5’-GATTATGCCTAATCTTGTTCTT-3’ and 5’-TAGCATTTTTCTTCCTCCTAAGAT-3’ was Dig labeled and used as a probe.

### Immunofluorescence assay

IFA was performed as described before (52) with few modifications. Briefly, 1 ml of parasite culture at 5% parasitemia was washed with PBS and re-suspended in a fresh fixative solution (4% Paraformaldehyde (EMS) and 0.0075% glutaraldehyde (EMS) in PBS). Fixed parasites were treated with 0.1% Triton-X 100 (Sigma) in PBS, then blocked with 3% BSA (Sigma) in PBS. Cells were then incubated with primary antibodies used at the following dilutions: mouse-anti-HA (Roche, 1:300), rabbit-anti-γ-H2A.X (Cell signaling, cat # 9718S, 1:300), rabbit-anti-Rad51 1:100 (Gene Tex cat # GTX100469) incubated for 1.5 h and washed three times in PBS. Samples were incubated with Alexa Fluor 488 goat anti-mouse (Life Technologies, 1:500) and Alexa Fluor 594 goat anti-rabbit (Life Technologies, 1:500). Samples were washed and laid on “PTFE” printed slides (EMS) and mounted in ProLong Gold antifade reagent with DAPI (Molecular Probes). Fluorescent images were obtained using a Plan Apo λ 100x oil NA=1.5 WD=130μm lens on a Nikon Eclipse Ti-E microscope equipped with a CoolSNAP Myo CCD camera. Images were processed using the NIS-Elements AR (4.40 version) software.

### Western blot

To collect parasite proteins, infected RBCs were lysed with saponin, then the parasites were washed twice with PBS and lysed in Laemmli sample buffer (Sigma). Equal amount of denatured protein samples were subjected to SDS-PAGE (gradient 4-20%, Bio-Rad) and electroblotted to a nitrocellulose membrane. Immunodetection was carried out using rabbit polyclonal anti-Halo antibody (Promega; 1:1000), mouse anti-HA antibody (Roche; 1:1000), rabbit-anti-γH2A.X antibody (Cell signaling cat # 9718S, 1:1000) and rabbit polyclonal anti-aldolase (1:2000) (51). The secondary antibodies used were antibodies conjugated to Horseradish Peroxidase (HRP), goat anti-rabbit and goat anti-mouse antibodies (Jackson ImmunoResearch Laboratories, 1:10000). The immunoblots were developed in EZ/ECL solution (Israel Biological Industries).

### X-ray irradiation of parasites

Parasites were irradiated using a PXi precision X-ray irradiator set at 225 kV, 13.28 mA. Prior to irradiation parasitemia was quantified by flow cytometry. A starting parasitemia of 0.5 % ring stage parasites in the control and the knockdown populations were both exposed to 10 & 60-Gy X-ray irradiation. After irradiation parasites were either collected immediately or put back in culture and media was replaced daily. Population recovery was measured by flow cytometry daily. To detect the increase in damage following irradiation, we collected parasites 15 minutes after irradiation and measured the levels of phosphorylated PfH2A by Western blot as described above.

### In Situ DNA Fragmentation (TUNEL) Assay

Tightly synchronized ring stages parasites (NF54 and *Pf*SR1) were fixed for 30 minutes in freshly prepared fixative (4% paraformaldehyde and 0.005 % glutaraldehyde). After fixation cells were rinsed three times with PBS and incubated with a permeabilization solution (0.1% Triton X-100 in PBS) for 10 min on ice. The cells were washed twice with PBS, and once with wash buffer supplied with the TUNEL Assay Kit-BrdU Red (Abcam cat # ab6610). Tunel assay was performed as per manufacturer guidelines. Briefly, following washing 50 μl of TUNEL reaction mixture (DNA labeling solution) was added to each sample. The cells were incubated for 60 min at 37°C with intermittent shaking. Cells were then washed three times with a rinse buffer (5 min each time) and re-suspended in 100 μl of antibody solution for 30 minutes at room temperature. Cells were then washed three times with PBS and mounted using Invitrogen™ Molecular Probes™ ProLong™ Gold Antifade reagent with DAPI, and imaged using a fluorescent microscope as described above.

### Co-Immunoprecipitaion

Immunoprecipitation was performed according to Grant S. Stewart et al., 2003 (53) with a slight modification. In brief, 200 ml of parasite cultures (∼10% parasitemia) were saponin lysed, and washed with PBS containing protease inhibitors. Subsequently, the parasite pellet was dissolved in chilled lysis buffer containing 50 mM Tris/HCl pH 7.5, 150 mM NaCl, 1 mM EDTA, 0.1% SDS and 1% NP40 supplemented with protease inhibitors (Roche) and sonicated for 4 cycles of 10-15 sec at 45% output using hielscher UP200S sonicator. The sonicated pellet was incubated for 30 minutes on ice. The lysate was purified by a few rounds of centrifugations at 10000x*g* for 10 min and incubated with a primary antibody anti-γ-H2A.X (Abcam cat # Ab2893, 1:300) for 10-12 h at 4°C with continuous swirling. The supernatant was further incubated for 12–14 h with Protein A/G agarose beads (Pierce) at 4°C, and then the beads were pelleted by centrifugation at 4°C. Beads were then washed with ice chilled washing buffer. Immunoprecipitated proteins were eluted with SDS Lamelli buffer and used for detection by SDS–PAGE and Western blot analysis.

### DNA repair assay

Tightly synchronized ring stage parasites were exposed to 10-Gy X-ray irradiation using a PXi irradiator as described above (19). Immediately following irradiation, parasites were put back to culture to allow them to repair the damaged DNA. Protein was extracted 15 min after irradiation (0h), as well as from parasite collected 3h and 6h after irradiation. Proteins extracted from untreated iRBCs were used as control. Western blot analysis was used to follow the changes in γ-PfH2A compared with the housekeeping control gene aldolase in each treatment. These Western blots were subjected to densitometry analysis to calculate the ratio between γ-PfH2A levels and aldolase.

## Acknowledgments

This work was supported partially by the Israeli Academy for Science, Israel Science Foundation (ISF) Grant 1523/18 and in part by European Research Council (erc.europa.eu) Consolidator Grant 615412 and Ministry of Science and Technology Grant 3-16285 (to R.D.). RD is also supported by the Dr. Louis M. Leland and Ruth M. Leland Chair in Infectious Diseases. BS and MG were supported by the PBC Fellowship Program for Outstanding Post-Doctoral Researchers from China and India. We also acknowledge Dr. Juan Lopez-Rubio and Dr. Dave Richard for kindly sharing plasmids that were used in this study.

## References

1. WHO. (2016) World Malaria Report, 2016.

2. Gardner, M.J., Hall, N., Fung, E., White, O., Berriman, M., Hyman, R.W., Carlton, J.M., Pain, A., Nelson, K.E., Bowman, S. et al. (2002) Genome sequence of the human malaria parasite Plasmodium falciparum. Nature, 419, 498–511.

3. Cartegni, L., Chew, S.L. and Krainer, A.R. (2002) Listening to silence and understanding nonsense: exonic mutations that affect splicing. Nat Rev Genet, 3, 285–298.

4. Eshar, S., Allemand, E., Sebag, A., Glaser, F., Muchardt, C., Mandel-Gutfreund, Y., Karni, R. and Dzikowski, R. (2012) A novel Plasmodium falciparum SR protein is an alternative splicing factor required for the parasites’ proliferation in human erythrocytes. Nucleic Acids Res, 40, 9903–9916.

5. Eshar, S., Altenhofen, L., Rabner, A., Ross, P., Fastman, Y., Mandel-Gutfreund, Y., Karni, R., Llinas, M. and Dzikowski, R. (2015) PfSR1 controls alternative splicing and steady-state RNA levels in Plasmodium falciparum through preferential recognition of specific RNA motifs. Mol Microbiol, 96, 1283–1297.

6. Shkreta, L. and Chabot, B. (2015) The RNA Splicing Response to DNA Damage. Biomolecules, 5, 2935–2977.

7. Naro, C., Bielli, P., Pagliarini, V. and Sette, C. (2015) The interplay between DNA damage response and RNA processing: the unexpected role of splicing factors as gatekeepers of genome stability. Frontiers in genetics, 6, 142.

8. Nathan, C. and Shiloh, M.U. (2000) Reactive oxygen and nitrogen intermediates in the relationship between mammalian hosts and microbial pathogens. Proc Natl Acad Sci U S A, 97, 8841–8848.

9. Gupta, D.K., Patra, A.T., Zhu, L., Gupta, A.P. and Bozdech, Z. (2016) DNA damage regulation and its role in drug-related phenotypes in the malaria parasites. Scientific reports, 6, 23603.

10. Kirkman, L.A., Lawrence, E.A. and Deitsch, K.W. (2014) Malaria parasites utilize both homologous recombination and alternative end joining pathways to maintain genome integrity. Nucleic Acids Res, 42, 370–379.

11. Singer, M., Marshall, J., Heiss, K., Mair, G.R., Grimm, D., Mueller, A.K. and Frischknecht, F. (2015) Zinc finger nuclease-based double-strand breaks attenuate malaria parasites and reveal rare microhomology-mediated end joining. Genome Biol, 16, 249.

12. Tarique, M., Ahmad, M., Chauhan, M. and Tuteja, R. (2017) Genome Wide In silico Analysis of the Mismatch Repair Components of Plasmodium falciparum and Their Comparison with Human Host. Front Microbiol, 8, 130.

13. Bethke, L., Thomas, S., Walker, K., Lakhia, R., Rangarajan, R. and Wirth, D. (2007) The role of DNA mismatch repair in generating genetic diversity and drug resistance in malaria parasites. Mol Biochem Parasitol, 155, 18–25.

14. Castellini, M.A., Buguliskis, J.S., Casta, L.J., Butz, C.E., Clark, A.B., Kunkel, T.A. and Taraschi, T.F. (2011) Malaria drug resistance is associated with defective DNA mismatch repair. Mol Biochem Parasitol, 177, 143–147.

15. Ahmad, M. and Tuteja, R. (2014) Emerging importance of mismatch repair components including UvrD helicase and their cross-talk with the development of drug resistance in malaria parasite. Mutat Res, 770, 54–60.

16. England, C.G., Luo, H. and Cai, W. (2015) HaloTag technology: a versatile platform for biomedical applications. Bioconjugate chemistry, 26, 975–986.

17. Prommana, P., Uthaipibull, C., Wongsombat, C., Kamchonwongpaisan, S., Yuthavong, Y., Knuepfer, E., Holder, A.A. and Shaw, P.J. (2013) Inducible knockdown of Plasmodium gene expression using the glmS ribozyme. PLoS One, 8, e73783.

18. Rogakou, E.P., Pilch, D.R., Orr, A.H., Ivanova, V.S. and Bonner, W.M. (1998) DNA double-stranded breaks induce histone H2AX phosphorylation on serine 139. J Biol Chem, 273, 5858–5868.

19. Goyal, M., Heinberg, A., Mitesser, V., Kandelis-Shalev, S., Singh, B.K. and Dzikowski, R. (2020) Phosphorylation of the canonical histone H2A marks foci of damaged DNA in malaria parasites. bioRxiv, 2020.2011.2006.372391.

20. Calhoun, S.F., Reed, J., Alexander, N., Mason, C.E., Deitsch, K.W. and Kirkman, L.A. (2017) Chromosome End Repair and Genome Stability in Plasmodium falciparum. mBio, 8.

21. Gopalakrishnan, A.M. and Kumar, N. (2015) Antimalarial action of artesunate involves DNA damage mediated by reactive oxygen species. Antimicrob Agents Chemother, 59, 317–325.

22. Ghorbal, M., Gorman, M., Macpherson, C.R., Martins, R.M., Scherf, A. and Lopez-Rubio, J.J. (2014) Genome editing in the human malaria parasite Plasmodium falciparum using the CRISPR-Cas9 system. Nat Biotechnol, 32, 819–821.

23. Dutertre, M., Lambert, S., Carreira, A., Amor-Gueret, M. and Vagner, S. (2014) DNA damage: RNA-binding proteins protect from near and far. Trends in biochemical sciences, 39, 141–149.

24. Chan, Y.A., Hieter, P. and Stirling, P.C. (2014) Mechanisms of genome instability induced by RNA-processing defects. Trends Genet, 30, 245–253.

25. Atamna, H. and Ginsburg, H. (1993) Origin of reactive oxygen species in erythrocytes infected with Plasmodium falciparum. Mol Biochem Parasitol, 61, 231–241.

26. Lee, A.H., Symington, L.S. and Fidock, D.A. (2014) DNA repair mechanisms and their biological roles in the malaria parasite Plasmodium falciparum. Microbiology and molecular biology reviews: MMBR, 78, 469–486.

27. Percario, S., Moreira, D.R., Gomes, B.A., Ferreira, M.E., Goncalves, A.C., Laurindo, P.S., Vilhena, T.C., Dolabela, M.F. and Green, M.D. (2012) Oxidative stress in malaria. International journal of molecular sciences, 13, 16346–16372.

28. Radfar, A., Diez, A. and Bautista, J.M. (2008) Chloroquine mediates specific proteome oxidative damage across the erythrocytic cycle of resistant Plasmodium falciparum. Free radical biology & medicine, 44, 2034–2042.

29. Tilley, L., Straimer, J., Gnadig, N.F., Ralph, S.A. and Fidock, D.A. (2016) Artemisinin Action and Resistance in Plasmodium falciparum. Trends Parasitol, 32, 682–696.

30. Mok, S., Ashley, E.A., Ferreira, P.E., Zhu, L., Lin, Z., Yeo, T., Chotivanich, K., Imwong, M., Pukrittayakamee, S., Dhorda, M. et al. (2015) Drug resistance. Population transcriptomics of human malaria parasites reveals the mechanism of artemisinin resistance. Science, 347, 431–435.

31. Li, P.C., Lam, E., Roos, W.P., Zdzienicka, M.Z., Kaina, B. and Efferth, T. (2008) Artesunate derived from traditional Chinese medicine induces DNA damage and repair. Cancer Res, 68, 4347–4351.

32. Berdelle, N., Nikolova, T., Quiros, S., Efferth, T. and Kaina, B. (2011) Artesunate induces oxidative DNA damage, sustained DNA double-strand breaks, and the ATM/ATR damage response in cancer cells. Mol Cancer Ther, 10, 2224–2233.

33. Kadioglu, O., Chan, A., Cong Ling Qiu, A., Wong, V.K.W., Colligs, V., Wecklein, S., Freund-Henni Rached, H., Efferth, T. and Hsiao, W.W. (2017) Artemisinin Derivatives Target Topoisomerase 1 and Cause DNA Damage in Silico and in Vitro. Front Pharmacol, 8, 711.

34. Leva, V., Giuliano, S., Bardoni, A., Camerini, S., Crescenzi, M., Lisa, A., Biamonti, G. and Montecucco, A. (2012) Phosphorylation of SRSF1 is modulated by replicational stress. Nucleic Acids Res, 40, 1106–1117.

35. Kornblihtt, A.R. (2007) Coupling transcription and alternative splicing. Adv Exp Med Biol, 623, 175–189.

36. Zhong, X.Y., Wang, P., Han, J., Rosenfeld, M.G. and Fu, X.D. (2009) SR proteins in vertical integration of gene expression from transcription to RNA processing to translation. Mol Cell, 35, 1–10.

37. Schor, I.E., Lleres, D., Risso, G.J., Pawellek, A., Ule, J., Lamond, A.I. and Kornblihtt, A.R. (2012) Perturbation of chromatin structure globally affects localization and recruitment of splicing factors. PLoS One, 7, e48084.

38. Loomis, R.J., Naoe, Y., Parker, J.B., Savic, V., Bozovsky, M.R., Macfarlan, T., Manley, J.L. and Chakravarti, D. (2009) Chromatin binding of SRp20 and ASF/SF2 and dissociation from mitotic chromosomes is modulated by histone H3 serine 10 phosphorylation. Mol Cell, 33, 450–461.

39. Sapra, A.K., Anko, M.L., Grishina, I., Lorenz, M., Pabis, M., Poser, I., Rollins, J., Weiland, E.M. and Neugebauer, K.M. (2009) SR protein family members display diverse activities in the formation of nascent and mature mRNPs in vivo. Mol Cell, 34, 179–190.

40. Ji, X., Zhou, Y., Pandit, S., Huang, J., Li, H., Lin, C.Y., Xiao, R., Burge, C.B. and Fu, X.D. (2013) SR proteins collaborate with 7SK and promoter-associated nascent RNA to release paused polymerase. Cell, 153, 855–868.

41. Sanford, J.R., Gray, N.K., Beckmann, K. and Caceres, J.F. (2004) A novel role for shuttling SR proteins in mRNA translation. Genes Dev, 18, 755–768.

42. Sanford, J.R., Ellis, J.D., Cazalla, D. and Caceres, J.F. (2005) Reversible phosphorylation differentially affects nuclear and cytoplasmic functions of splicing factor 2/alternative splicing factor. Proc Natl Acad Sci U S A, 102, 15042–15047.

43. Michlewski, G., Sanford, J.R. and Caceres, J.F. (2008) The splicing factor SF2/ASF regulates translation initiation by enhancing phosphorylation of 4E-BP1. Mol Cell, 30, 179–189.

44. Karni, R., Hippo, Y., Lowe, S.W. and Krainer, A.R. (2008) The splicing-factor oncoprotein SF2/ASF activates mTORC1. Proc Natl Acad Sci U S A, 105, 15323–15327.

45. Fregoso, O.I., Das, S., Akerman, M. and Krainer, A.R. (2013) Splicing-factor oncoprotein SRSF1 stabilizes p53 via RPL5 and induces cellular senescence. Mol Cell, 50, 56–66.

46. Aley, S.B., Sherwood, J.A. and Howard, R.J. (1984) Knob-positive and knob-negative Plasmodium falciparum differ in expression of a strain-specific malarial antigen on the surface of infected erythrocytes. J Exp Med, 160, 1585–1590.

47. Calderwood, M.S., Gannoun-Zaki, L., Wellems, T.E. and Deitsch, K.W. (2003) Plasmodium falciparum var genes are regulated by two regions with separate promoters, one upstream of the coding region and a second within the intron. J Biol Chem, 278, 34125–34132.

48. Theriault, C. and Richard, D. (2017) Characterization of a putative Plasmodium falciparum SAC1 phosphoinositide-phosphatase homologue potentially required for survival during the asexual erythrocytic stages. Scientific reports, 7, 12710.

49. Deitsch, K., Driskill, C. and Wellems, T. (2001) Transformation of malaria parasites by the spontaneous uptake and expression of DNA from human erythrocytes. Nucleic Acids Res, 29, 850–853.

50. Sambrook, J., Fritsch, E. and Maniatis, T. (1989) Molecular Cloning: A Laboratory Manual. Cold Spring Harbor Laboratory, New York.

51. Dahan-Pasternak, N., Nasereddin, A., Kolevzon, N., Pe’er, M., Wong, W., Shinder, V., Turnbull, L., Whitchurch, C.B., Elbaum, M., Gilberger, T.W. et al. (2013) PfSec13 is an unusual chromatin associated nucleoporin of Plasmodium falciparum, which is essential for parasite proliferation in human erythrocytes. J Cell Sci.

52. Dahan-Pasternak, N., Nasereddin, A., Kolevzon, N., Pe’er, M., Wong, W., Shinder, V., Turnbull, L., Whitchurch, C.B., Elbaum, M. and Gilberger, T.W. (2013) PfSec13 is an unusual chromatin-associated nucleoporin of Plasmodium falciparum that is essential for parasite proliferation in human erythrocytes. J Cell Sci, 126, 3055–3069.

53. Stewart, G.S., Wang, B., Bignell, C.R., Taylor, A.M. and Elledge, S.J. (2003) MDC1 is a mediator of the mammalian DNA damage checkpoint. Nature, 421, 961–966.

